# Mapping microglia and astrocytes activation *in vivo* using diffusion MRI

**DOI:** 10.1101/2020.02.07.938910

**Authors:** Raquel Garcia-Hernandez, Antonio Cerdán Cerdá, Alejandro Trouve Carpena, Mark Drakesmith, Kristin Koller, Derek K. Jones, Santiago Canals, Silvia De Santis

**Author notes:** Corresponding: Silvia De Santis,; Santiago Canals.

## Abstract

Glia, and particularly microglia, are increasingly implicated in the pathophysiology of psychiatric and neurodegenerative disorders. However, to date the only methods for imaging these cells in vivo involve either invasive procedures (e.g. multi-photon imaging in rodents) or TSPO-PET radiotracers, which afford low resolution and specificity, since TSPO expresses across multiple cell types. Here, we present a non-invasive diffusion-weighted MRI method to image changes in glia morphometry *in vivo*. Using two rat models of neuroinflammation, with and without neurodegeneration, we demonstrate that diffusion-weighted MRI carries the fingerprint of microglia and astrocytes activation, and that specific signatures from each population can be quantified non-invasively. We demonstrate that the method can further detect glia proliferation, and provide a quantitative account of neuroinflammation regardless of the existence of a concomitant neuronal loss. We prove the translational value of the approach showing significant correlations between MRI and histological microglia markers measured across different brain regions in humans. This framework holds the potential to transform basic and clinical research by providing a tool to clarify the role of inflammation in health and disease across the lifespan.

## Introduction

Neurodegenerative diseases such as Alzheimer’s, Parkinson’s, multiple sclerosis and dementia are a pressing problem for developed societies with aging populations (*1-3*). Accumulating evidence suggests chronic neuroinflammation, the sustained activation of microglia and astrocytes, to strongly influence neurodegeneration and contribute to its progression. A major question is whether inhibition of the inflammatory response has the ability to reverse or slow down its symptoms (*4*). In addition, abnormal immune activation during puberty and adolescence has been associated with increased vulnerability to brain disorders later in life (*5*), making the characterization of the inflammatory profile along the lifespan a hot topic. Therapies targeting glial cells are currently being proposed as disease-altering treatments to improve the outcome of neurological disorders, with extremely promising results (*4,6-7*). Furthermore, for many brain diseases, neuroinflammation is emerging as a cause, rather than a consequence of the pathogenesis (*8*); thus, characterizing the tissue inflammatory state could serve as valuable early disease biomarkers. In this context, desired properties of such biomarkers would be the capacity to detect both changes in morphology and proliferation/depletion (both hallmarks of glia activation), and, importantly, to discriminate inflammation with and without neurodegeneration.

While imaging techniques are widely adopted to monitor neurologic conditions, non-invasive approaches able to specifically characterize brain inflammation *in vivo* are lacking. The current gold-standard is positron emission tomography (PET)-based targeting of the 18kDa translocator protein. While difficult to generalize due to different binding genotypes across individuals (*9*), PET is associated with ionizing radiation exposure, which limits its use in vulnerable populations and longitudinal studies, and also has low spatial resolution, making it unsuitable to image small structures. In addition, inflammation-specific radiotracers express across multiple cell types (microglia, astrocytes and endothelium). Lastly, a significant tracer uptake in the periphery makes it hard to separate central from peripheral inflammation (*10*). Diffusion-weighted magnetic resonance imaging (DW-MRI), on the other hand, has the unique ability to image brain microstructure *in vivo*, non-invasively and with high resolution by capturing the random motion of water molecules in brain parenchyma (*11*).

While the DW-MRI signal is potentially sensitive to all extracellular and intracellular spaces that restrict water displacement in the tissue, current formulations are mostly designed for white matter and axons. A few recent studies, establishing the groundwork for this work, showed that conventional MRI signal can be sensitive to various alterations in microglia (*12-14*), but none so far showed specificity to microglia and astrocyte activation, or inflammation in presence of neurodegeneration. Achieving specificity is of key importance as neurodegenerative diseases manifest through different mechanisms, involving specific cell populations, all playing potentially different roles in disease causation and progression. Importantly, by combining advanced DW-MRI sequences with mathematical models based on neurobiological knowledge of brain parenchyma morphology, the diffusion characteristics within specific tissue compartments, and even cell types, could be measured.

With this idea in mind, we developed an innovative strategy to image microglia and astrocyte activation in grey matter using DW-MRI, by building a microstructural multi-compartment tissue model informed by knowledge of microglia and astrocyte morphology. To validate the model, we first used an established rat paradigm of inflammation based on intracerebral lipopolysaccharide (LPS) administration (*15*). In this paradigm, neuronal viability and morphology is preserved, while inducing microglial activation within few hours, and a delayed astrocytic response which is detectable only 24 hours after injection (*16*). Therefore, glial responses can be transiently dissociated from neuronal degeneration, and the signature of reactive microglia investigated independently of any astrogliosis. Then, to isolated the imaging fingerprint of astrocyte activation, we repeated the LPS experiment but pretreating the animals with the CSF1R-inhibitor PLX5622 (Plexxikon Inc.), which is known to temporally deplete around 90% of the microglia (*17*). Finally, we used an established paradigm of neuronal damage, based on ibotenic acid administration (*18*), to test whether the model was able to disentangle neuroinflammatory signatures with and without a concomitant neurodegeneration. This is pivotal for demonstrating the utility of the framework as a biomarker discovery platform for the inflammatory status in neurodegenerative diseases, where both glia activation and neuronal damage are key players.

We demonstrate, for the first time, that DW-MRI signal carries the fingerprint of microglial and astrocyte activation, with signatures specific to each glia population, reflecting the morphological changes as validated *post-mortem* by quantitative immunohistochemistry. Importantly, we demonstrate that the framework is both sensitive and specific to inflammation with and without neurodegeneration, so that the two conditions can be teased apart. In addition, we demonstrate the translational value of the approach in a cohort of healthy humans, in which we performed a reproducibility analysis. Significant correlation with known microglia density patterns in the human brain supports the utility of the method to generate reliable glia biomarkers. A framework able to characterize relevant aspects of tissue microstructure during inflammation, *in vivo* and non-invasively, is expected to have a tremendous impact on our understanding of the pathophysiology of many brain conditions, and transform current diagnostic practice and treatment monitor strategies.

## Materials and Methods

### Animal preparation

All animal experiments were approved by the Animal Care and Use Committee of the Instituto de Neurociencias de Alicante, Alicante, Spain, and comply with the Spanish (law 32/2007) and European regulations (EU directive 86/609, EU decree 2001-486, and EU recommendation 2007/526/EC). Rats were housed in groups (4-5), with 12-12h light/dark cycle, lights on at 8:00, at room temperature (23 ± 2°C) and free access to food and water. Glia activation was achieved by intracranial injection in the dorsal hippocampus (coordinates bregma -3.8 mm, sup-inf 3.0 mm, 2 mm from midline in the left hemisphere) of 2 μl of saline and LPS at concentration of 2.5 μg/μl. The opposite hemisphere was injected with the same amount of saline. In a cohort of animals, microglia depletion was achieved by administering the CSF1R-inhibitor PLX5622 (Plexxikon Inc.) in two ways: as a dietary supplement in standard chow at 1200 ppm (Research Diets), and with intraperitoneal injection of 50 mg/kg in vehicle once a day for 7 days. Another cohort of rats received the same chow without enrichment, and was injected IP once a day with the same doses of vehicle. Less than 24 hours after the last injection, all the rats were injected LPS according to the procedure described above. Neuronal death was achieved injecting 1 μl of saline and ibotenic acid at concentration of 2.5 μg/μl in the dorsal hippocampus (same coordinates). The opposite hemisphere was injected with the same amount of saline. A subgroup of animals underwent mynocycline treatment (45 mg/Kg; Sigma-Aldrich, Madrid, Spain) according to (*18*). Mynocycline was administered intraperitoneally (i.p) 12 h prior to the surgery, 30 min before the surgery and once a day for three days at 24 h intervals.

After different post-injection delays, the rats were scanned in the MRI scanner and immediately perfused for *ex-vivo* MRI immunohistological analysis of microglia (Iba-1+) and astrocytes (GFAP+). A total of 36 rats were used, with weights in the range 250gr-300gr, divided in 5 groups. Group 1 (n=6) received the LPS injection, and was scanned and perfused after 8 hours. Group 2a (n=7) received the LPS injection and was scanned and perfused after 24 hours. Group 2b (n=4) was treated with control chow and injected with vehicle for 7 days, then received the LPS injection and was scanned and perfused after 24 hours. Group 3 (n=6) was treated with PLX5622 for 7 days, then received the LPS injection and was scanned and perfused after 24 hours. Group 4 (n=6) received the LPS injection and was scanned and perfused after a minimum of 15 days, or more if the ventricular enlargement was not reabsorbed. No statistically significant differences were detected between group 2a and b (control for PLX5622 chow), so they were merged into a single group for the rest of the analysis. Group 5a and 5b (n=9) received ibotenic acid injection and was scanned and perfused after 14 days post-surgery. Group 5b (n=6) was treated with Mynocycline for 5 days.

Experimental design is reported in Fig. S1.

### MRI experiment

#### Rats

MRI experiments on rats were performed on a 7 T scanner (Bruker, BioSpect 70/30, Ettlingen, Germany) using a receive-only phase array coil with integrated combiner and preamplifier in combination with an actively detuned transmit-only resonator. DW-MRI data were acquired using an EchoPlanar Imaging diffusion sequence, with 30 uniform distributed gradient directions, b = 2000 and 4000 s/mm2, diffusion times 15, 25, 40 and 60 ms with four non-diffusion weighted images, repetition time (TR) = 7000 ms and echo time (TE) = 25 ms. Fourteen horizontal slices were set up centered in the hippocampus with field of view (FOV) = 25 × 25 mm^2^, matrix size = 110 × 110, in-plane resolution = 0.225 × 0.225 mm^2^ and slice thickness = 0.6 mm. Additionally, three relaxometry sequences were acquired with the same geometry of the DW-MRI scan: a gradient echo sequence with TR = 1500 ms, 30 TE equally spaced between 3.3 and 83.4 ms and 3 averages; a T1-weighted sequence with TR = 300 ms. TE = 12.6 ms and 2 averages; and a T1-weighted sequence with TR = 3000 ms. TE = 7.7 ms and 4 averages. Finally, a high-resolution anatomical scan with full brain coverage was acquired with TR = 8000 ms. TE = 14 ms, 4 averages, FOV = 25 × 25 mm^2^, matrix size = 200 × 200, in-plane resolution = 0.125 × 0.125 mm^2^, 56 slices of thickness = 0.5 mm. Total scan time including animal positioning was around 2 hours.

#### Humans

6 healthy subjects were scanned 5 times in a 3T Siemens Connectom scanner. DW-MRI data were acquired using an EchoPlanar Imaging diffusion sequence with the following parameters: TE=80 ms, TR 3.9 s, diffusion times 17.3, 30, 42 and 55 ms, bvalues 2000 and 4000 s/mm2, each with 30 and 60 uniformly orientated gradient directions, respectively, and 6 unweighted (b=0) images per diffusion time, yielding a total 384 images. Additional parameters used were: flip angle 90, slice thickness 2mm, in-plane voxel size 2mm, field of view 220×220 mm, matrix size 110×110. Total scan time was around 40 minutes.

### MRI analysis and statistics

Rat MRI data were processed as follows. Preliminary data were used to verify the reach of a 2μl injection, confirming that the liquid filled the whole dentate gyrus. So, regions of interest (ROIs) for the analysis were manually drawn in the dentate gyrus of the dorsal hippocampus; the injection tract was used to locate the central slice, and ROIs were drawn from two slices before the injection, and up to 2 slices after, for a total length of 3 mm covered.

Raw DW-MRI data were non-linearly registered to the T2-weighted scan to correct for EPI distortions, corrected for motion and eddy current distortions using affine registration, then fed to a custom routine written in Matlab (R2018a, the Mathworks) which fits the signal to a multi-compartment model (MCM) of diffusion The MCM is inspired by the AxCaliber model for white matter (*31*) but is adapted to grey matter morphology. The model comprises: one compartment of water undergoing restricted diffusion in cylindric geometry (representing water trapped into cell ramifications) with a main orientation and a Watson dispersion term (*26*), two spherically restricted compartments (*32*), one extracellular space matrix, aligned with the main cylinder orientation and modelled as a tensor, and one free water compartment, modelled according to (*21, 26*). In formulas:

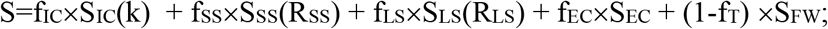

Where f_IC_ is the fraction of water undergoing restricted diffusion in cylinders, S_IC_ is the signal in Watson-dispersed cylinders expressed according to eq. 2 of (*26*), k is the Watson dispersion parameter, f_SS_ and f_LS_ are the fractions of water undergoing restricted diffusion in the two spherical compartments, S_SS_ and S_LS_ are the signals of water undergoing restricted diffusion in spheres expressed according to eq. 15 of (*32*), which depend on the radii of the two spherical compartments R_SS_ and R_LS_, respectively, f_EC_ is the fraction of water hindered in the extracellular space, S_IC_ is the signal in the extracellular space modeled as a tensor with radial symmetry, whose main orientation is linked to the main orientation of the cylindrical compartment, 1-f_T_ is the fraction of free water (defined as one minus the tissue fraction f_T_ to help the discussion), and S_FW_ is the free water signal, defined as in (*21*). The fitting parameters are f_IC_, f_SS_ and f_LS_, k, R_SS_ and R_LS_, the extra-cellular tensor diffusivity and f_T_. The MCM is illustrated in Fig. S2.

The low b-value shell was used to fit the conventional tensor model and produce maps of the mean diffusivity. T1 and T2 weighted maps were also used to calculate the T1/T2 ratio, which is considered a proxy for myelination (*33*). T2* maps were calculating by fitting an exponential decay to the T2*-weighted images acquired at different echo times. Additionally, for illustration purposes the high-resolution anatomical scans were non-linearly registered to a rat brain template using an advanced normalization approach (*35*). Repeated measure ANOVA was used to check for significant effect of the injection and of the group. Post-hoc t-tests were used to compare injected versus control hemisphere, and corrected for multiple comparisons.

Similarly, human MRI data were preprocessed as follows. Motion, eddy current and EPI distortions were corrected using FSL TOPUP and EDDY tools (*36*). Correction for gradient non-linearities (*37-38*), signal drift (*39*) and Gibbs ringing artefacts (*40*) was also performed. All diffusion data were then registered to a skull-stripped (*41*) structural T1-weighted image using EPIREG (*36*).

B0 scans were non-linearly registered to a high-resolution human brain template *(42*) using an advanced normalization approach (*35*); then, the inverse transformation was applied to bring the Desikan GM parcellation in the single subject space. Mean values of all MRI parameters were calculated by averaging the maps in the left and right hippocampi, the cerebellum, frontal, occipital and motor cortex. Both intra and inter-subject coefficient of variations were calculated for each MRI measure. The Desikan parcellization was used to calculate mean and standard deviation of the stick fraction in 7 regions of interest, that were correlated with post-mortem histological staining for microglia density, as reported in (*23*).

### Tissue processing and immunohistochemistry

Rats were deeply anesthetized with a lethal dose of sodium pentobarbital, 46mg/kg, injected intraperitoneally (Dolethal, E.V.S.A. laboratories., Madrid, España). Rats were then perfused intracardially with 100ml of 0.9% phosphate saline buffer (PBS) and 100ml of ice-cold 4% paraformaldehyde (PFA, BDH, Prolabo, VWR International, Lovaina, Belgium). Then, brains were immediately extracted from the skull and fixed for 1 hour in 4% PFA. Afterwards, brains were included in 3% Agarose/PBS (Sigma-Aldrich, Madrid, Spain), and cut in vibratome (VT 1000S, Leica, Wetzlar, Germany) in 50 μm thick serial coronal sections.

Coronal sections were rinsed and permeabilized three times in 1xPBS with Triton X-100 at 0.5% (Sigma-Aldrich, Madrid, Spain) for 10 minutes each. Then, they were blocked in the same solution with 4% of bovine serum albumin (Sigma-Aldrich, Madrid, Spain) and 2% of goat serum donor herd (Sigma-Aldrich, Madrid, Spain) for 2 hours at room temperature. The slices were then incubated for one night at 4°C with primary antibodies for Iba-1 (1:1000, Wako Chemicals, Osaka, Japan), GFAP (1:1000, Sigma-Aldrich, Madrid, Spain) and neurofilament 360Kd medium (1:250, Abcam, Cambridge, United Kingdom) and NeuN (1:250, Sigma-Aldrich, Madrid, Spain) to label microglia, astrocytes, neuron processes and nuclei respectively. The sections were subsequently incubated in specific secondary antibodies conjugated to the fluorescent probes, each at 1:500 (ThermoFisher Scientific, Waltham, USA) for 2h at room temperature. Sections were then treated with 4′,6-Diamidine-2′-phenylindole dihydrochloride at 15mM (DAPI, Sigma-Aldrich, Madrid, Spain) during 15 minutes at room temperature. Finally, sections were mounted on slides and covered with an anti-fading medium using a mix solution 1:10 Propyl-gallate:Mowiol (P3130, SIGMA-Aldrich, Madrid, Spain; 475904, MERCK-Millipore, Massachussets, United States).

For myelin labelling, antigen retrieval was performed in 1% citrate buffer (Sigma-Aldrich, Madrid, Spain) and 0.05% of Tween 20 (Sigma-Aldrich, Madrid, Spain) warmed to 80° for protein unmasking. The rest of the steps were done as described above, using myelin basic protein primary antibody (MERCK-Millipore, Massachussets, United States).

#### Imaging and data extraction

The tissue sections were then examined using a computer-assisted morphometry system consisting of a Leica DM4000 fluoresce microscope equipped with a QICAM Qimaging camera 22577 (Biocompare, San Francisco, USA) and Neurolucida morphometric software (MBF, Biosciences, VT, USA). Microglia were visualized and reconstructed under Leica HC PLC APO objective 20x/0.5 and astrocytes under Leica HC PLC APO objective 40x/0.75. Five cells per hippocampus per hemisphere were randomly selected for a total of 720 cells included for analysis (360 microglia, 360 astrocytes). Only cells that displayed intact and clear processes were included. Cells were traced through the entire thickness of the sections, and trace information was then saved as 3D reconstructions or rendered into a 2-dimensional diagram of each cells following analysis requirement.

Metric analysis of reconstructed cells was extracted using Neurolucida Explorer software (MBF, Biosciences, VT, USA) and Imaris (Bitplane, Belfast, United Kingdom): cell body perimeter, number of primary processes, number of nodes (branch points), complexity ([Sum of the terminal orders + Number of terminals] * [Total dendritic length / Number of primary dendrites]), fibres density and dendograms, cell size and polar plots (Fig. S5). Polar plots were analyzed in order to extract fiber orientation and the dispersion parameter in a plane parallel to the microscope, where higher values mean more uniform distribution of the fibers around the cell body. Importantly, 3D convex analysis was performed to estimate astrocytes volume and overcome the limitations of GFAP labelling (*43-45*). The volume estimation in the analysis is defined as the area of the polygon created from straight lines connecting the most distal points of the astrocytes processes.

Density analysis was performed on 12-bit grey scale pictures acquired with the described system. ROIs were manually delineated following the Franklin and Paxinos rat brain atlas (*34*), covering the complete hippocampus in each hemisphere, for at least 5 slices per rat. Analysis was performed using Icy software (*45*) in a semi-automatic manner. The threshold for detection of positive nuclei was set for each condition, setting average nuclei size and a signal/noise ratio higher than 23%, according to Rayleigh criterion for resolution and discrimination between two points.

Myelin, neurofilament and neural nuclei fluorescent analysis was also performed on pictures acquired with the described system and analyzed using Icy Software (*45*). Two ROIs of 200 µm^2^ were placed per hippocampus per hemisphere in at least 5 slices per rat to obtains the corresponding intensity values.

#### Data analysis and statistics

The statistical analysis was done using GraphPad Prism 7 software (GraphPad Software Inc., La Jolla, CA, USA) and Rstudio (RStudio 2015. Inc., Boston, MA). The presence of outlier values and parametric distribution was checked. We applied unpaired t-test for comparing, for each time point, control hemisphere versus injected hemisphere. Pearson’s correlation was used for regression analysis, and coefficients were transformed to apply Fisher’s 1925 test (*46*) for significant values. Polar analysis and dispersion estimation were performed to obtain the dispersion parameter (*47*).

## Results

### Microstructural model of diffusion and immunological challenge in rats

We built a model of grey matter diffusivity, as detailed in the methods section. Briefly, the model accounts for water diffusion in the microglial compartment corresponding to small cell somas (modeled as small spheres) with thin cellular processes (modeled as sticks) growing radially with a degree of dispersion, and an astrocytic compartment consisting of large globular cells (modeled as large spheres) (*19*). It is important to note that while GFAP stains the cytoskeleton of the cell, astrocytes have a globular shape (*20*). Both compartments are embedded in an extracellular space compartment, composed by a tensor-like sub-compartment (hindered water in contact with structures and cells) and a free water compartment of water undergoing unrestricted diffusion. Importantly, changes in the tissue fraction (1 minus the free water signal), is a surrogate measure of tissue loss, i.e., of degeneration (*21*). We then tested the mathematical model with two experimental paradigms. In the first one, neuroinflammation without neurodegenerations is induced by an injection of LPS, and in the second, neuroinflammation with degeneration is induced by an injection of ibotenic acid. All injections were performed in one hemisphere (targeting the dentate gyrus of the hippocampus), with the contralateral side as within-subjects control (vehicle injected, see methods for details).

### Microglia activation characterized using Iba-1 staining and MRI

Morphological analysis of Iba-1^+^ cells by histology in the tissue at different time-points after LPS injection demonstrated a fast microglial reaction with retraction of cellular processes at 8h, that progressed at 24 h with an additional increase in the microglial cell body size and an increase in the process dispersion parameter (indicating less dispersion), as shown in Fig. 1 b-c. No changes in cell density are found (Fig. S6). The distinct and time-dependent changes in microglial cell morphology were tightly mirrored by the imaging parameters specifically related to the microglia compartment, i.e., sticks (cellular processes) and the small spheres (cell soma). Accordingly, the stick fraction was significantly reduced at 8 hours in the injected versus control hippocampus, and further reduced at 24 hours. Conversely, the radius of the small sphere component was significantly increased at 8 and 24 hours. Importantly, these LPS-induced changes disappeared when the animals were depleted of microglia by pretreatment with PLX5622, demonstrating the specificity of these MRI signatures to microglia. Moreover, when looking at inter-individual variability, a strong association was found between MRI-derived microglial markers and their histological counterparts at all measured time points (Fig. 1e-f). Finally, at 2 week post-injection, when complete recovery was expected, all parameters measured from both histology and MRI converged, showing no statistically significant difference between injected and control hemisphere. Overall, these results demonstrated the possibility to recover a microglia-specific signal from DW-MRI with capacity to unmask a microglial reaction *in vivo* (Fig. 1g).

**Fig. 1:**
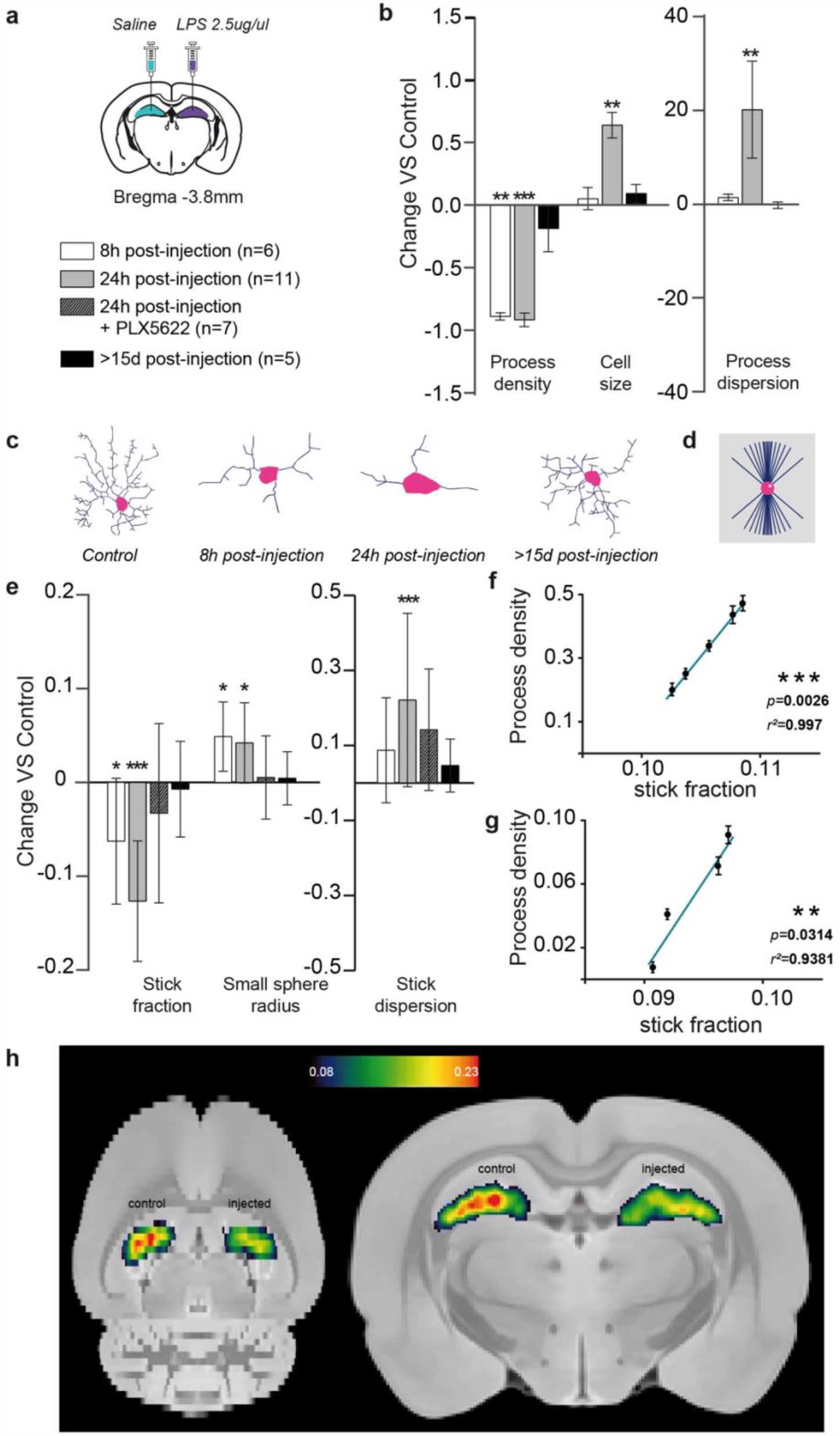
Histological characterization of microglia reaction and its associated MR imaging signature. **a**, experimental scheme showing bilateral stereotaxic injection of LPS (left) / saline (right) and the composition of the four groups: 6 animals were scanned 8 hours post-injection, 11 animals were scanned 24 hours post-injection, 8 animals were treated with PLX5622 for 7 days before the injection and then scanned 24 hours post-injection, and 5 animals were scanned 15 days or more post-injection. **b**, normalized change (P_injected_-P_control_)/P_control_ in process density, cell size and process dispersion factor for the injected versus control hippocampus, measured in Iba-1^+^ stained microglia for the different groups. Asterisks represent significant paired difference between injected and control. Error bars represent standard deviation. **c**, morphology reconstruction of representative microglia at the different times. **d**, geometrical model used for microglia. **e**, normalized change (P_injected_-P_control_)/P_control_ between MRI parameter calculated in the injected vs control hemisphere for the microglia compartment. Asterisks represent significant paired difference between injected and control. **f, g**, correlations between stick fraction from MRI and process density from Iba-1 at 8 (e) and 24 hours post injection (f). **h**, mean stick fraction maps at 24 h post-injection, normalized to a rat brain template and averaged over all rats.

### Astrocytes activation characterized using GFAP staining and MRI

We next performed a comparable analysis with astrocytes (labeled as GFAP^+^ cells) taking advantage of the distinct time course of their response to LPS injection. This cell population, unlike microglia, showed no significant alteration in either density or morphology at 8h post LPS injection, as shown in Fig. 2b and supplementary Fig 7. However, at 24h, astrocytes grow in volume as measured by the mean radius of the convex hull (Fig. 2, see methods for details). Interestingly, the associated MRI compartment for astrocytes, i.e. the large spheres, followed the same pattern of changes across conditions. Volume of GFAP^+^ cells and the mean radius of the large spheres measured by DW-MRI grew in parallel at 24h post LPS injection, were insensitive to microglia depletion with PLX5622 and recovered towards baseline levels at 15 days post-injection. Accordingly, their inter-individual variability showed a strong correlation (Fig. 2d). Therefore, the results obtained for the astrocytic component also demonstrate the possibility of recovering an astrocyte-specific signal from DW-MRI and the capacity to map astrocytic reactions *in vivo* (Fig. 1g).

**Fig. 2:**
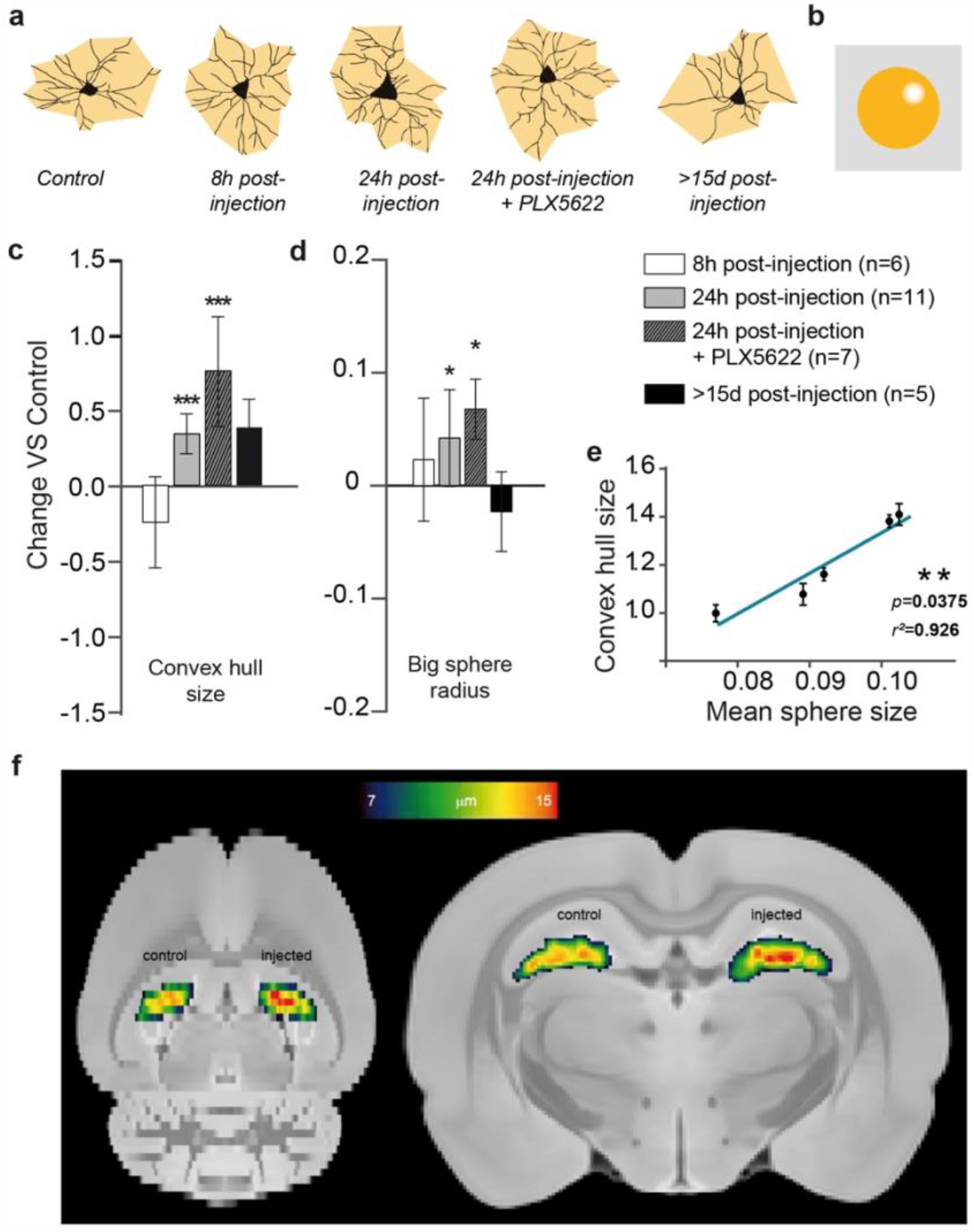
Histological characterization of astrocyte reaction and its associated MR imaging signature. **a**, morphology reconstruction of representative astrocytes at the different times in black and 2-D convex hull in grey. **b**, geometrical model used for astrocites. **c**, normalized change (P_injected_-P_control_)/P_control_ in convex hull mean radius for the injected versus control hippocampus, measured in GFAP^+^ stained astrocytes for the different groups. Asterisks represent significant paired difference between injected and control. Error bars represent standard deviation. **d**, normalized change (P_injected_-P_control_)/P_control_ between MRI-derived large sphere radius calculated in the injected vs control hemisphere for the astrocyte compartment (shown in the insert). Asterisks represent significant paired difference between injected and control. **e**, correlation between mean sphere radius from MRI and convex hull mean radius from GFAP. **f**, large sphere radius maps at 24 h post-injection, normalized to a rat brain template and averaged over all rats.

### Concomitant microglia activation and neuronal death characterized using NeuN staining and MRI

To challenge the capability of the developed model to distinguish between pure inflammation and inflammation with concomitant neurodegeneration, a cohort of animals was injected as before, but with ibotenic acid, using the contralateral (right) hemisphere as control (saline injected). Histological staining demonstrated that ibotenic acid at the chosen concentration induced a microglial reaction characterized by retraction of the cellular processes with increased dispersion index, and a significant increase in cell density (Fig. 3 a-b). No alterations in astrocytes were observed (data not shown). Neuronal staining with NeuN unveiled a large decrease in staining intensity in the injected hemisphere, demonstrating the severe neuronal loss induced by ibotenic acid (*14b*). These histological findings were tightly mirrored by the MRI parameters (Fig. 3 c), with a significant decrease of stick fraction, increase in the stick dispersion parameter and, notably, increase in the small sphere density. In Fig. 3 c, the MRI parameters measured in the LPS cohort at 8h post-injection, where we detected microglia activation but no neuronal damage (Fig. 1), are shown here in white to facilitate the comparison of hallmarks of a glia reaction with and without neuronal damage. Furthermore, a distinct signature for microglia proliferation, captured by the small sphere density, differentiating LPS from ibotenic lesions, could be extracted from the DW-MRI data (Fig. 3c).

**Fig. 3:**
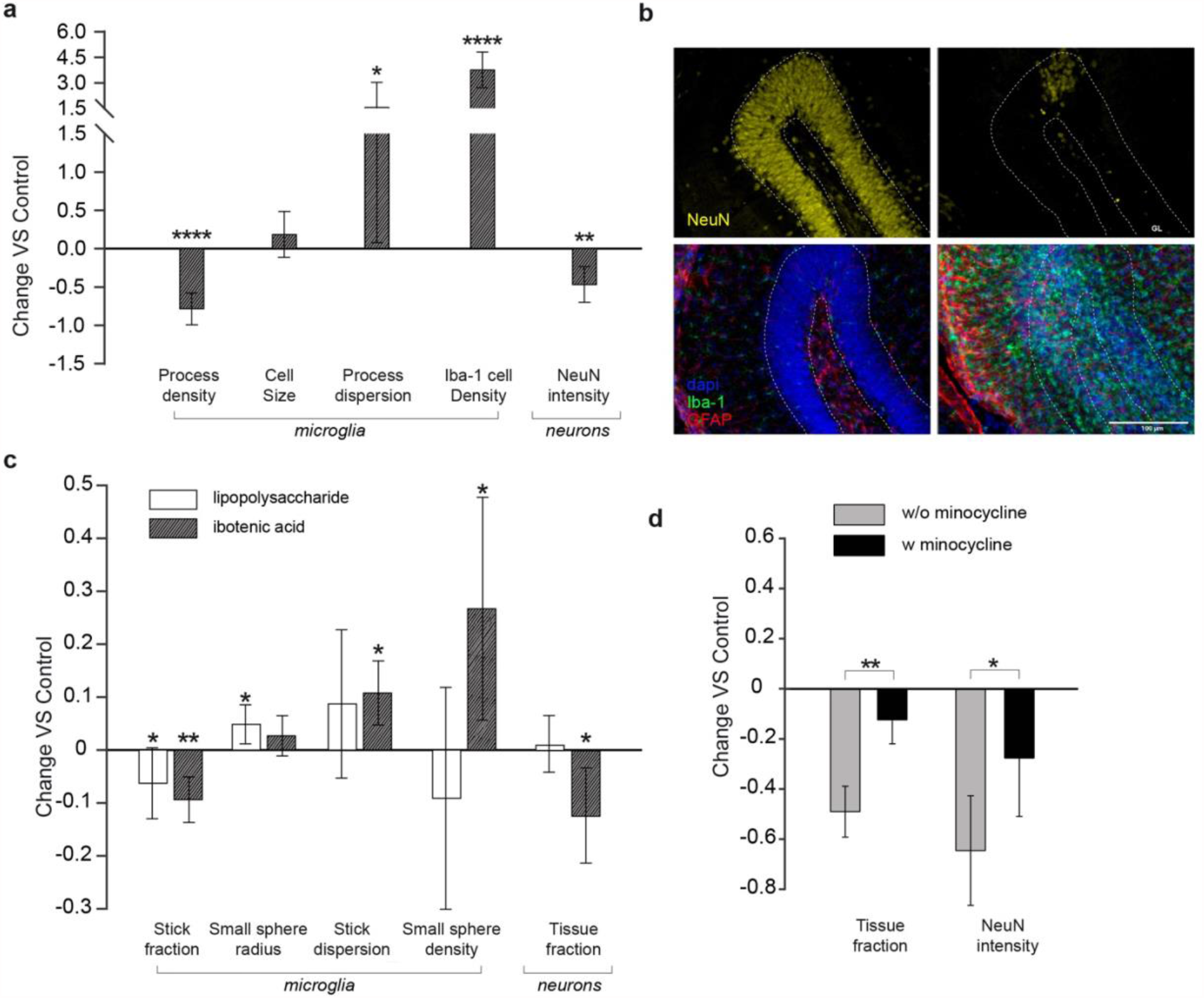
Characterization of inflammation in the presence of neuronal death. **a**, normalized change (P_injected_-P_control_)/P_control_ in histological measures for the injected versus control hippocampus. Asterisks represent significant paired difference between injected and control. **b**, NeuN and GFAP-Iba1 staining of a representative animal (left, control; right, injected). **c**, normalized change (P_injected_-P_control_)/P_control_ between MRI parameter calculated in the ibotenic-injected vs control hemisphere for microglia and neuron compartments (dark grey). For comparison, the same parameters obtained in group 2 of the LPS-injected animals is reported in white. Asterisks represent significant paired difference between injected and control. **d**, normalized change (P_injected_-P_control_)/P_control_ for MRI and histological markers of neuronal death calculated separately in the untreated animals, and in those treated with minocycline. Asterisks represent significant unpaired difference between the two groups.

Interestingly, the tissue fraction component was significantly decreased in the injected hemisphere, compared to control, and only for the ibotenic acid injection not for LPS, suggesting an association with neuronal degeneration. To test this hypothesis, we pretreat the animals with minocycline, an anti-inflammatory drug (*18*), and repeated the ibotenic acid injections as before. NeuN staining demonstrated the protective effect of minocycline on ibotenic-induced neuronal death and, importantly, the MRI parameter tissue fraction captured this effect, showing a significant reduction of the ibotenic-induced decrease in this parameter. These results demonstrated the utility of the tissue fraction component to monitor neuronal degeneration.

### Comparison with conventional MRI techniques

To highlight the importance of the developed framework, it is important to show that conventional MRI is sensitive to morphological changes due to inflammation, but cannot disentangle the different populations involved across different conditions, as shown in Fig. S4. Glia activation caused an increase in mean diffusivity, but different stages could not be differentiated. A clear reduction of T1/T2 is observed in all three conditions, while histology demonstrates no myelin change (Fig. S3). T2* does not have enough sensitivity to reflect glia morphological changes at any stages.

### Translation to human

As a proof of concept for the translational validity of these results and to evaluate the reproducibility of the proposed imaging framework, we adapted the MRI protocol to a human 3T Connectom scanner (*22*) and acquired data from a healthy cohort. As shown in Fig. 4, the multi-compartment model applied to these data returned values for the within-subjects coefficient of variations (CoV) in the range 1.5-8%, and between-subjects CoV in the range 2.6-15%, which are in the range of conventional MRI measures routinely used in the clinics with diagnostic value. Finally, we took advantage of the known heterogeneous distribution of microglial cell densities across brain regions in humans to test the ability of our framework to quantify microglial cell populations *in vivo*. Importantly, we found that the stick fraction calculated across different grey matter regions follows the patterns of microglia cell density measured *post mortem* in humans (*23*), with a significant correlation between histological and MRI measures (p=0.03, Fig. 5).

**Fig. 4:**
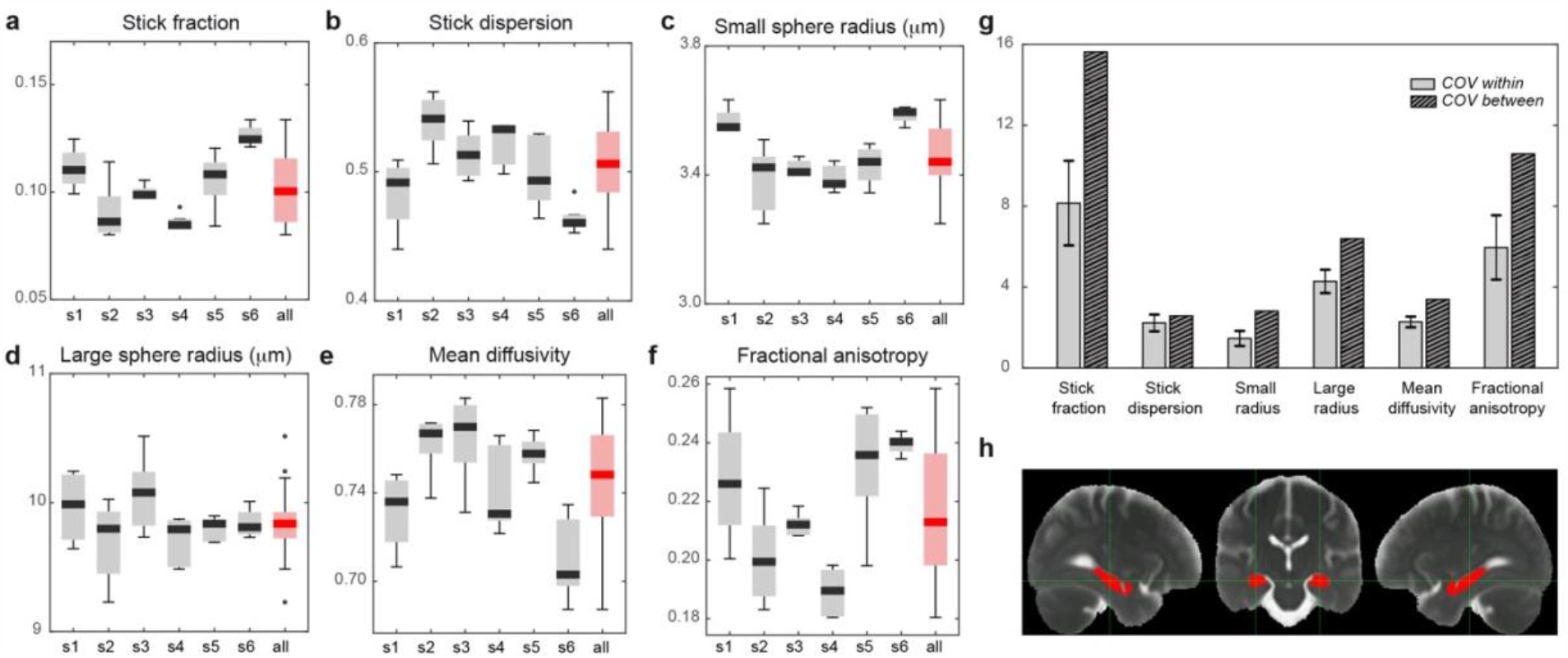
Feasibility of the framework translation to humans and MR parameters reproducibility analysis. **a**, boxplot of stick fraction as measured separately in the hippocampus of 6 subjects scanned 5 times (s1-s6) and pooling all subject together (red). The same is shown for the stick dispersion (**b**), small (**c**), large sphere radius (**d**), mean diffusivity (**e**) and fractional anisotropy (**f**). **g**, average coefficient of variation calculated within subject (light grey) and between subjects (striped). **h**, region of interest in the hippocampus used for the reproducibility analysis, defined according to (*42*).

**Fig. 5:**
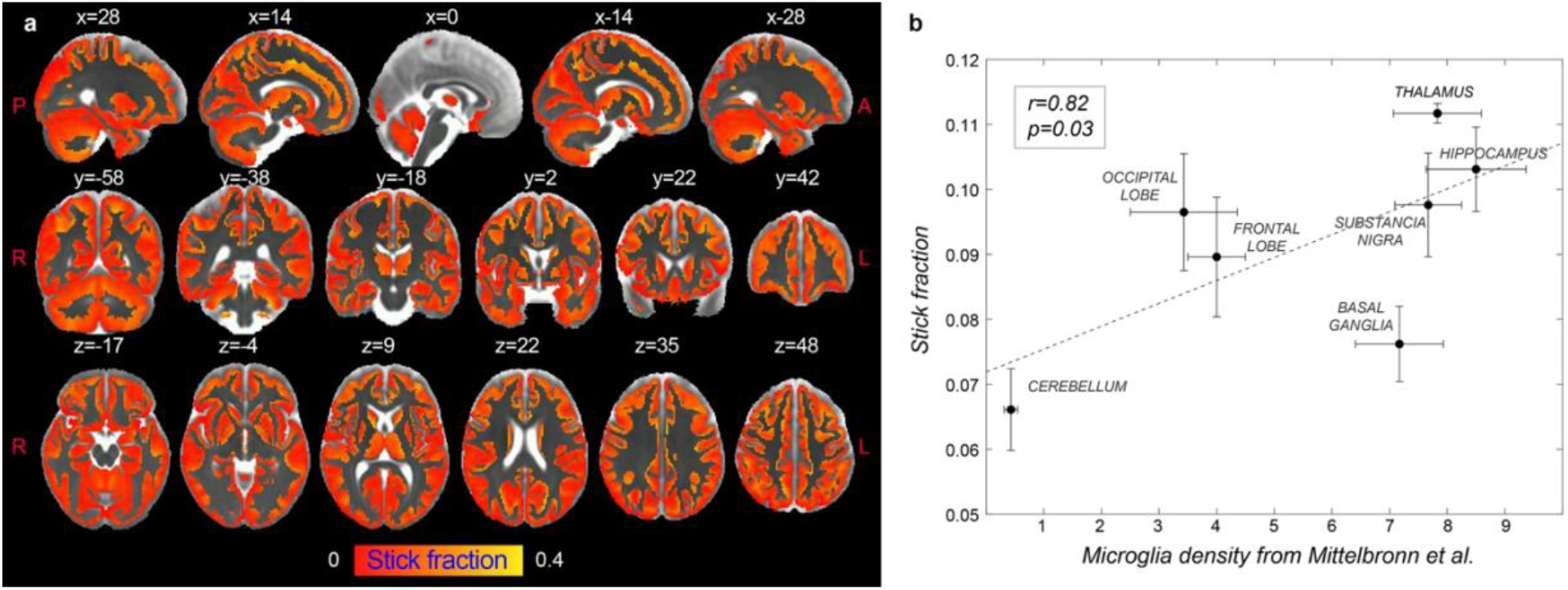
Correlation between the stick fraction and microglia density in human brain. **a**, stick fraction according to the multi-compartment model normalized to the brain template defined in (*42*), masked for grey matter tissue, and averaged across subjects. **b**, correlation between stick fraction calculated in 7 grey matter regions (hippocampus, cerebellum, substancia nigra, thalamus, motor, frontal and occipital cortex) and microglia density measured using histological staining of post-mortem human tissue as reported in (*23*). Error bars represent standard deviation.

## Discussion

Diffusion MRI signal has great potential to reveal the inflammatory component in numerous brain conditions (*24*), and several efforts have been made to provide microstructural models able to capture features belonging to distinct tissue sub-compartments, for example by including dendrite dispersion (*25, 26*) or a compartment for the soma of neurons (*27*). However, to date, no framework is available to specifically look at the cellular signature of glia activation. Here we propose and validate a strategy to image microglia and astrocyte activation in grey matter using diffusion MRI, and demonstrate its translational validity to humans. By taking advantage of the different activation windows of glia in an LPS-driven immunological challenge in rats, and by using pharmacological tools to deplete microglia in the brain, we were able to dissect the MRI signatures of specific glial responses. We identified two MRI parameters, namely the stick fraction and the larger sphere size, as sensitive to, and only to, microglia and astrocyte activation, respectively. In addition, using injections of ibotenic acid we demonstrate that the framework is able to distinguish glia activation and proliferation, independently of an underlying neurodegenerative processes. Glia proliferation is an important aspect of the inflammatory reaction and a major component in the evolution of chronic neurodegeneration (*28*). Modulating ibotenic acid-induced neurodegeneration with anti-inflammatory pretreatments, we further unveil an MRI parameter with capacity to monitor neuronal loss. The possibility of teasing apart the contribution of inflammation and neuronal loss has direct implications for understanding the contribution of the brain innate immune response to disease progression, where both components are key players in the pathophysiology and can be targeted by disease-modifying treatments.

Our results are supported by quantitative cell morphology analysis. Validation of MRI results is challenging due to several factors, including the need to co-register regions of interest with very different sizes and properties and the need for tissue fixation in histological preparations (*29*). To overcome this limitation, here we relied on measuring changes rather than absolute magnitude of quantities. We demonstrate that there is a very high correlation between the changes found in injected *vs*. control regions, as measured using MRI and histology, suggesting that our imaging measures truly capture the hallmark of glia activation with high sensitivity.

The proposed MRI methodology was adapted to a human MRI scanner, and healthy subjects were recruited to perform a reproducibility study, demonstrating that the glia biomarkers are highly reproducible between different MRI sessions and in line with CoVs calculated for conventional MRI parameters routinely used in clinical settings (*30*). Importantly, *in vivo* variability of MRI-derived microglia density follows known patterns of cell density measured *post-mortem* in humans, with significant correlations between the stick fraction and microglia density calculated in several regions of the brain parenchyma.

Our results have implications in the interpretation of several imaging studies published so far. On one hand, we propose that since diffusion MRI signal is sensitive to glia activation, the biological substrate of some of the alterations reported in numerous brain conditions could be driven by changes in glia morphology, rather than the conventional interpretation as “neural damage or degeneration”. For example, multiple sclerosis (MS) causes not only demyelination and neuronal damage, but also inflammation (*2*), which is likely contributing to the observed differences in MRI parameters between control and patients. On the other hand, our results obtained with traditional MRI, in agreement with recently published literature (*12-14*), show that conventional parameters are sensitive to morphological changes due to inflammation, but cannot disentangle the different populations involved (microglia, astrocytes, neurons). This implies that the biological substrate of the observed changes is invisible to conventional MRI.

This study has some limitations. While the multicompartment model does not explicitly differentiate between microglia processes and neuronal dendrites, basic geometrical reasoning supports the idea that axons and dendrites reorganization would impact the stick dispersion in a different way than microglia process retraction, and would thus be easily distinguished from a microglia activation, although this claim needs to be proven in future studies. However, the MRI results obtained in the animal cohort which received injection of ibotenic acid demonstrated that the proposed framework is already capable of distinguishing between glia activation with and without neuronal loss, the fundamental pathological feature of neurodegenerative diseases.

To conclude, we proposed here a new generation of non-invasive glia-centric biomarkers, which are expected to transform the study of many diseases associated with a glial response: those where inflammation is as a known or possible cause, as well as those in which the glial reaction can serve as a powerful early diagnostic and/or prognostic marker.

## Notes

### Competing Interest Statement

The authors have declared no competing interest.

### Summary of Updates

Including neurodegeneration with ibotenic acid injections

